# Segregating domain-general from emotional context-specific inhibitory control systems - ventral striatum and orbitofrontal cortex serve as emotion-cognition integration hubs

**DOI:** 10.1101/2020.12.09.418830

**Authors:** Qian Zhuang, Lei Xu, Feng Zhou, Shuxia Yao, Xiaoxiao Zheng, Xinqi Zhou, Jialin Li, Xiaolei Xu, Meina Fu, Keshuang Li, Deniz Vatansever, Keith M. Kendrick, Benjamin Becker

## Abstract

Inhibitory control hierarchically regulates cognitive and emotional systems in the service of adaptive goal-directed behavior across changing task demands and environments. While previous studies convergently determined the contribution of prefrontal-striatal systems to general inhibitory control, findings on the specific circuits that mediate the context-specific impact of inhibitory control remained inconclusive. Against this background we employed an evaluated emotional Go/No Go task with fMRI in a large cohort of subjects (N = 250) to segregate brain systems and circuits that mediate domain-general from emotion-specific inhibition control. Particularly during a positive emotional context, behavioral results showed a lower accuracy for No Go trials and a faster response time for Go trials. While the dorsal striatum and lateral frontal regions were involved in inhibitory control irrespective of emotional context, activity in the ventral striatum (VS) and medial orbitofrontal cortex (mOFC) varied as a function of emotional context. On the voxel-wise network level, limbic and striatal systems generally exhibited highest changes in global brain connectivity during inhibitory control, while global brain connectivity of the left mOFC was less suppressed during emotional contexts. Functional connectivity analyses moreover revealed that negative coupling between the VS with inferior frontal gyrus (IFG)/insula and mOFC varied as a function of emotional context. Together these findings indicated separable domain general systems as well emotional context-specific inhibitory brain systems which specifically encompass the VS and its connections with frontal regions.

## 1. Introduction

Cognitive control refers to the hierarchical regulation of other cognitive and emotional systems in the service of adaptive goal-directed behavior. The inhibition of habitual or prepotent behavioral responses represents a cardinal cognitive control domain and is essential for adaptive regulation of behavior across changing task demands and environments. Performance in this domain is influenced by personal and contextual factors. Response inhibition has been associated with traits such as impulsivity (Dalley and Robbins, 2017) and deficient response inhibition has been reported across major psychiatric disorders, including drug addiction (Courtney et al., 2013; Lüscher et al., 2020), depression (Diener et al., 2012; Liu et al., 2020; Wessa et al., 2007), schizophrenia (Ursu et al., 2011; Vercammen et al., 2012) and attention deficit/hyperactivity spectrum disorders (ADHD, Karalunas et al., 2020). Response inhibition is also modulated by the emotional context, as revealed by a growing number of behavioral findings using affectively charged task stimuli in Go/No Go paradigms (Albert et al., 2012; Drevets and Raichle, 1998; Hare et al., 2005; Putman et al., 2010). A number of studies reported longer reaction times for negative stimuli and more commission errors (failed suppression of the response) for positive stimuli (Albert et al., 2012; Hare et al., 2005; Putman et al., 2010; Schulz et al., 2007), while others reported no modulatory effects (Albert et al., 2010; Brown et al., 2012).

Animal models and human neuroimaging studies have provided convergent evidence for a pivotal role of the striatum and regulatory frontal regions in response inhibition and suggest that their functional interplay may be critical for adaptive inhibition (Hu et al., 2019; Lüscher et al., 2020; Morein-Zamir and Robbins, 2015; Volkow et al., 2011). Across species the underlying neural systems have been mapped primarily in the domain of motor-response inhibition by employing the Go/No Go paradigm that assesses the suppression of a prepotent motor response before its initiation and the stop-signal reaction time task (SSRT) that measures suppression after the motor response has been initiated. Animal models that focused on the specific contribution of striatal subregions indicate that the inhibition of prepotent motor responses is mediated by the dorsal striatal systems (Eagle et al., 2008, 2011; Eagle and Baunez, 2010) and human neuroimaging studies employing translational paradigms confirm the role of the dorsal striatum (DS) in motor inhibition (Ghahremani et al., 2012; Kelly et al., 2004; Robertson et al., 2015) while additionally emphasizing the contribution of the anterior cingulate and lateral prefrontal regions, particularly the inferior frontal gyrus (IFG) (Aron et al., 2004; Brown et al., 2012; Cromheeke and Mueller, 2014; Meffert et al., 2016; Shafritz et al., 2006; Swick et al., 2008). With respect to differentiating specific regions and circuits that mediate the influence of emotional context on response inhibition animal models are currently lacking and previous neuroimaging studies in humans revealed inconsistent results. Whereas a number of neuroimaging studies reported emotional context-specific recruitment of different regions including the striatal, amygdala-hippocampal and medial orbitofrontal regions during inhibitory control (Berkman et al., 2009; Goldstein et al., 2007; Hare et al., 2005; Protopopescu et al., 2005; Roberts et al., 2013; Somerville et al., 2011; Todd et al., 2012; Torrisi et al., 2016; Wessa et al., 2007; Zhang et al., 2016), subsequent meta-analyses have revealed rather inconclusive results, with emotional context-dependent recruitment of the anterior cingulate and precuneus (Cromheeke and Mueller, 2014), the amygdala (Hung et al., 2018) or a domain-general fronto-parietal network for emotional and cognitive control being reported (Chen et al., 2018; Xu et al., 2016). These diverging findings with respect to specific brain regions and circuits mediating the influence of emotional context on inhibitory control may be (partly) explained by the comparably low sample size or the use of a priori defined regions of interest in the original studies as well as the varying selection criteria for paradigm classes in the meta-analyses.

To separate brain regions and circuits that specifically mediate domain-general and emotional context-specific cognitive control the present study employed a validated emotional linguistic Go/No Go fMRI paradigm during which a large cohort of healthy subjects (N = 250) were required to inhibit prepotent motor responses in neutral, positive and negative contexts. In addition to task-based activation changes accumulating evidence highlights the contribution of network-level interactions during distinct cognitive functions such as working memory (Vatansever et al., 2015, 2017) or response inhibition (Tsvetanov et al., 2018). To facilitate a hypothesis-free determination of the corresponding network-level mechanism that underlies domain-general versus emotional-context specific cognitive control a data-driven network-level approach (intrinsic connectivity contrast, ICC) was employed that operates on the voxel-level and independent of pre-specified regions of interest. We initially validated the modulatory influence on the behavioral level and in line with previous studies expected that subjects would tend to exhibit longer reaction times for negative stimuli and make more errors during the suppression of responses during positive stimuli (Albert et al., 2012; Hare et al., 2005; Liu et al., 2020; Putman et al., 2010; Schulz et al., 2007). Based on previous animal models and human neuroimaging studies we expected that dorsal striatal and lateral prefrontal regions would exhibit a general engagement during inhibitory control, while regions engaged in emotional and valence processing, specifically the amygdala, ventral striatum and orbitofrontal regions (Goldstein et al., 2007; Hare et al., 2005; Taylor et al., 2018) would exhibit emotional context-specific engagement during inhibitory control. Given the pivotal role of the fronto-striatal cicruits in inhibitory control and their sensitivity to emotional context (Chang et al., 2020; Christakou et al., 2004; Li and Sinha, 2008; Somerville et al., 2011), we further hypothesized that core regions of this circuitry would exhibit emotional context-sensitive modulations on the network level.

## 2. Materials and Methods

### 2.1 Participants

N = 250 healthy, right-handed Chinese students were enrolled in the present study after providing written informed consent. Exclusion criteria included: (1) past or current psychiatric or neurological disorder, and (2) regular or current use of licit and illicit drugs or medication. Data from n = 11 subjects were lost due to technical failure (n = 5, MRI system failure, n = 6, failure of the behavioral data acquisition system). Data from n = 12 subjects were excluded due to excessive head motion during MRI acquisition (>2.5mm or 2.5 degrees). A total of n = 227 subjects were included in the final behavioral and fMRI analyses (112 males; age: mean ± SD = 21.62 ± 2.32 years). The study was approved by the local ethics committee and in accordance with the latest revision of the Declaration of Helsinki.

### 2.2 Emotional Go/No Go fMRI Paradigm

Cognitive control and the emotional modulation of cognitive control was examined by means of a modified version of a previously validated mixed event-related block design emotional (linguistic) Go/No Go paradigm (Goldstein et al., 2007; Protopopescu et al., 2005). Briefly, the paradigm uses orthographical cues to indicate the required behavioral response. Participants were instructed to silently read words and to respond as quickly and accurately as they can to the font of the word. For stimuli presented in the normal front (Go trials), participants were required to respond via right index finger button-press but to inhibit their response to words presented in italicized front (No Go trials). Instruction was presented for 1400ms before each block. Emotional context was modulated by presenting neutral, positive and negative stimuli. The stimuli consisted of 18 neutral, 18 negative and 18 positive words written in Chinese and were matched for length (length = 4) and word frequency. An independent sample of n = 18 subjects rated the emotional category (yes or no), intensity (1-9 scale), and imaginability (1-9 scale) of the word stimuli before the start of the study (details see **Supplementary Table S1**). The paradigm encompassed 24 blocks that were presented in 2 runs with 12 blocks each. Each run comprised two blocks of the six permutations of the response (Go, No Go) and emotion (positive, negative, neutral) conditions: neutral Go (neu Go), neutral No Go (neu No Go), negative Go (neg Go), negative No Go (neg No Go), positive Go (pos Go), positive No Go (pos No Go). During each go block, 18 words in normal font (100% Go trials) were presented while during no-go blocks 12 normal font words (66.7% Go trials) and 6 italicized font words (33.3% No Go trials) were presented.

Stimuli were presented for 300ms, followed by an inter-stimulus-interval (ITI) of 900ms, leading to a block duration of 21.6s. Blocks were interspersed by a low-level inter-block period of 16s. The blocks and trials were presented in pseudo-random and counter-balanced order (same block or trial never presented in succession). Total duration of the paradigm was approx. 16 minutes (**Figure 1**).

**Figure 1.**
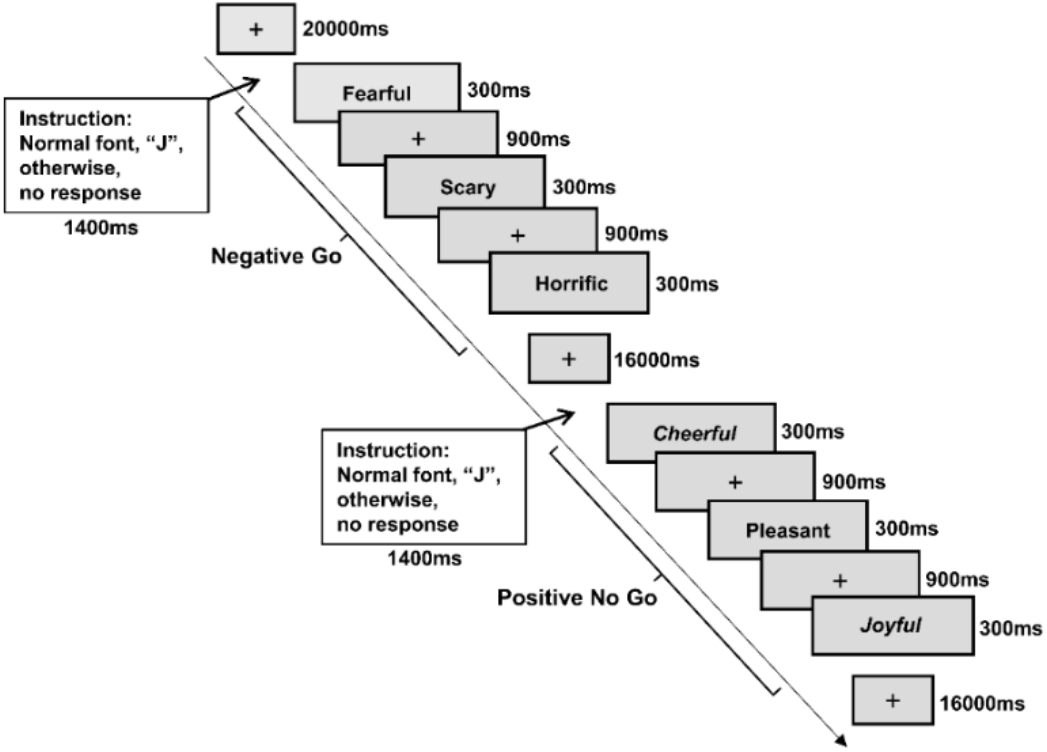
Experimental paradigm and timing. A negative Go block and positive No Go block are depicted for the emotional linguistic Go/No Go task.

### 2.3 Behavioral Data Analysis

Accuracy and response times (in Go trials) served as indices to examine the effect of inhibitory control and emotional context on the behavioral level and were examined by means of repeated measures ANOVA models. For accuracy, a two-way repeated-measures ANOVA with inhibition (Go vs. No Go) and emotion (negative vs. neutral vs. positive) as within-subject factors was employed. For response time in Go trials (correct trials), a one-way repeated measures ANOVA with the different emotional contexts was performed (negative vs. neutral vs. positive). For all behavioral analyses appropriate Bonferroni corrections were employed to account for multiple comparisons.

### 2.4 MRI Data Acquisition

MRI data were collected using a 3 Tesla, GE Discovery MR750 system (General Electric Medical System, Milwaukee, WI). For each subject, 488 volumes of T2*-weighted echo planar images were obtained using the following acquisition parameters: repetition time, 2000ms; echo time, 30 ms; slices, 39; slice-thickness, 3.4mm; gap, 0.6mm; field of view, 240 × 240 mm^2^; matrix size, 64 × 64; flip angle, 90°. To improve normalization of the functional images high-resolution whole brain T1-weighted images were acquired (spoiled gradient echo pulse sequence with oblique acquisition, acquisition parameters: repetition time, 6ms; echo time, minimum; flip angle, 9°; field of view =256 × 256mm; acquisition matrix, 256 × 256; thickness, 1mm; 156 slices). OptoActive MRI headphones (http://www.optoacoustics.com/) were used to reduce acoustic noise exposure during scanning.

### 2.5 MRI Data Analysis

#### 2.5.1 fMRI Data Preprocessing

MRI data were analyzed using SPM12 software (Wellcome Trust Center of Neuroimaging, University College London, London, United Kingdom). The first 10 volumes were excluded to allow active noise canceling and to gain magnet-steady data. The remaining time-series were slice timing corrected, realigned to correct for head movement and co-registered to the T1-weighted structural images. For normalization the T1-weighted images were segmented and the corresponding transformation matrices were applied to normalize the functional time-series to Montreal Neurological Institute (MNI) standard space. Normalized data were resampled to 3mm^3^ and smoothed using a Gaussian kernel with full-width at half-maximum (FWHM) of 8mm.

#### 2.5.2 BOLD Level Analysis

The first level analyses employed an event-related analysis approaches as implemented in the general linear model. Six condition-specific regressors (neu Go, neu No Go, neg Go, neg No Go, pos Go, pos No Go) were modelled on the first level and convolved with the standard hemodynamic response function (HRF). The six head-motion parameters were included as nuisance regressors to further control for movement-related artifacts. Contrast images of interest (all No Go > all GO, neu No Go > neu Go, neg No Go > neg Go, pos No Go > pos Go, No Go > Go) were created on the individual level. To examine general cognitive control networks and the modulation by emotional context, the following second level model were computed: (1) a one sample t test on all No Go > Go contrasts, corresponding to a main effect of emotion-unspecific inhibition, and (2) one-way repeated measures ANOVA including the emotion-specific contrasts of interest (neu No Go > neu Go, neg No Go > neg Go, pos No Go > pos Go), corresponding to an inhibition x emotional context interaction effect.

#### 2.5.3 Functional Connectivity Analysis (Intrinsic Connectivity Contrast)

To determine connectivity profiles during inhibitory control and their modulation by emotional context an intrinsic connectivity contrast (ICC) analysis was implemented. Initially, a strict noise regression was conducted for the preprocessed data using the SPM12-based Conn functional connectivity toolbox (Whitfield-Gabrieli and Nieto-Castanon, 2012) to remove noise from white matter, cerebrospinal fluid, six head-motion parameters and the second-order derivatives. In addition, a linear detrending and a temporal band-pass filter of 0.008 Hz - inferior was applied.

Next, six condition-specific regressors (neu Go, neu No Go, neg Go, neg No Go, pos Go, pos No Go) were modelled and voxel-based global connectivity (ICC maps) were calculated on each condition for each participant. The strength of the connectivity of each voxel was determined by the average r^2^ value (Martuzzi et al., 2011). The main effect of inhibition and inhibition x emotion interaction effects were determined using identical second-level models as employed for the activation analysis.

#### 2.5.4 Follow-up seed-to-whole brain connectivity analyses

To further determine the specific pathways involved in the emotional modulation of inhibitory control seed-to-voxel wise context-dependent functional connectivity analyses were computed using gPPI (gPPI) (McLaren et al., 2012). Given the hypothesized role of the striatal-frontal circuits in the emotional modulation of inhibitory control (Chang et al., 2020; Li and Sinha, 2008; Somerville et al., 2011), the analyses focused on identified regions within this circuitry. Specifically, for the follow-up functional connectivity analyses seeds (4-mm spheres) were placed at the peak coordinates of the inhibition x emotion interaction effects from the BOLD level (ventral striatum) and ICC analysis (mOFC). Main effects of inhibition and inhibition x emotion interaction effects were examined using one sample t-tests (contrast: No Go > Go) and one-way ANOVAs (contrasts: neu No Go > neu Go, neg No Go > neg Go, pos No Go > pos Go).

#### 2.5.5 Thresholding and regions of interest

The main aim of the BOLD level analyses was to segregate regions exhibiting a general involvement in inhibitory control from regions that exhibit an emotional modulation. To facilitate a clear separation a conservative peak-level threshold was applied on the whole brain level (p < 0.05 family-wise error, FWE) and only clusters > 10 voxels are reported. To allow a more sensitive examination on the network level a cluster-level correction approach was applied for the whole-brain ICC and functional connectivity analyses (p < 0.05 FWE, cluster level correction, initial cluster defining threshold of p < 0.001). The striatum-focused analysis for the ICC follow-up functional connectivity analyses aimed at determining the specific regions that exhibit an emotional modulation within the striatum and thus employed a small volume correction approach using a mask encompassing the entire striatum based on the Human Brainnetome Atlas (Fan et al., 2016) in combination with peak-level FWE corrected at p < 0.05. To further disentangle the direction of the emotion-specific effects, parameter estimates were extracted from 4-mm spheres centered on the peak voxels of significant interaction effects and subjected to post-hoc analyses.

## 3. Results

### 3.1 Behavioral Results

To examine the effect of emotional context on inhibitory control at the behavioral level, the repeated-measures ANOVA on the accuracy data revealed significant main effects of inhibition and emotional context as well as a significant inhibition x emotion interaction effect. Examination of the significant main effect of inhibition (F(1, 226) = 747.84, p < 0.001, η_p_^2^ = 0.77), revealed a higher accuracy for Go as compared to No Go trials (Go trials: mean ± SD = 98.54% ± 3.11%, No Go trials: mean ± SD = 70.81% ± 16.19%, Cohen’s d = 2.38). Examining the significant main effect of emotion (F(2, 452) = 6.64, p = 0.002, η_p_^2^ = 0.029) revealed a generally lower accuracy in the positive condition compared to both, the negative (positive: mean ± SD = 83.89% ± 10.08%, negative: mean ± SD = 85.24% ± 9.26%, p = 0.004, Cohen’s d = 0.14) as well as neutral condition (positive: mean ± SD = 83.89% ± 10.08%, neutral: mean ± SD = 84.90% ± 8.89%, p = 0.026, Cohen’s d = 0.11). Post-hoc analyses of the significant interaction effect (F(2, 452) = 4.22, p = 0.016, η_p_^2^ = 0.018) demonstrated that for No Go trials accuracy was significantly lower in the positive condition compared to both the negative (positive: mean ± SD = 69.40% ± 18.43%, negative: mean ± SD = 71.81% ± 17.36%, p = 0.01, Cohen’s d = 0.13) and neutral condition (positive: mean ± SD = 69.40% ± 18.43%, neutral: mean ± SD = 71.24% ± 16.54%, p = 0.046, Cohen’s d = 0.11, **Figure 2A**).

**Figure 2.**
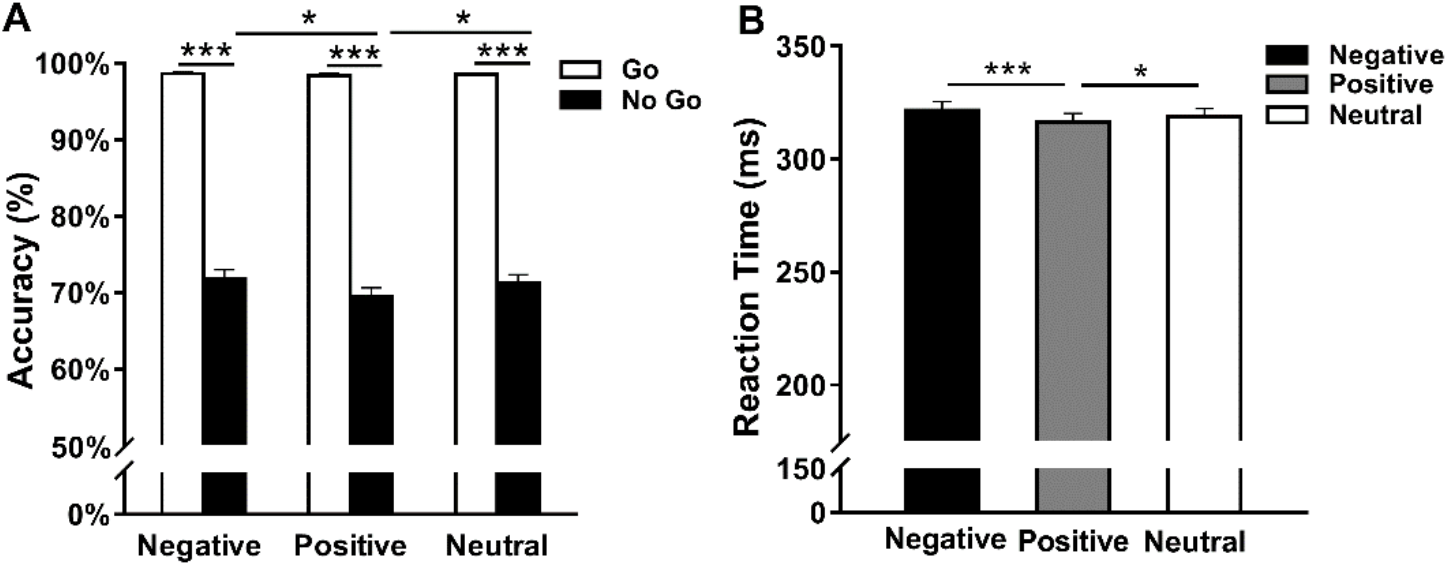
Response accuracy and reaction time in different emotional contexts. (A) Accuracy for Go trials and No Go trials across different emotional conditions. (B) Reaction time for Go correct trials across different emotional conditions. Bar graphs show mean ± SEM. * and *** denote significant post-hoc difference at p_Bonferroni_ < 0.05 and p_Bonferroni_ < 0.001 respectively.

For response times on Go trials, one-way repeated measures ANOVA revealed a main effect of emotion (F(2, 452) = 10.76, p < 0.001, η_p_^2^ = 0.045) with post-hoc tests demonstrating faster response times during the positive condition compared to both, the negative (positive: mean ± SD = 316.45 ± 55.95, negative: mean ± SD = 321.49 ± 58.43, p < 0.001, Cohen’s d = 0.09) as well as neutral condition (positive: mean ± SD = 316.45 ± 55.95, neutral: mean ± SD = 318.93 ± 57.60, p = 0.038, Cohen’s d = 0.04, **Figure 2B**).

### 3.2 fMRI Results

#### 3.2.1 BOLD Level Analysis

Examining the inhibitory control networks across emotional contexts (contrasts: No Go > Go) revealed widespread activity in fronto-parietal regions, primarily in the dorsolateral prefrontal (DLPFC), inferior lateral frontal and anterior insular (AI) cortex and superior parietal regions as well as striatal, primarily dorsal striatal regions (**Figure 3A and Table 1**). The whole-brain ANOVA examining the emotional modulation of inhibition (contrasts: neu No Go > neu Go, neg No Go > neg Go, pos No Go > pos Go) revealed a significant interaction effect mainly located in the bilateral posterior IFG/anterior insula (AI) (left, Z = 6.13, p_FWE_ < 0.001, x/y/z: - 39/15/-12; right, Z = 7.45, p_FWE_ < 0.001, x/y/z: 51/18/-3) as well as mOFC (left, Z = Inf, p_FWE_ < 0.001, x/y/z: −15/33/-15; right, Z = Inf, p_FWE_ < 0.001, x/y/z: 15/33/-12) and striatal regions, with additional clusters located in temporal, occipital, parietal, frontal and thalamic regions (details see **Supplementary Table S2 and Figure 3B**). Notably, in contrast to the main effect that mainly engaged the dorsal striatum the interaction effect was primarily located in bilateral ventral striatum (left, Z = 6.04, p_FWE_ = 0.002, voxels = 38, x/y/z: −12/18/-9; right, Z = 5.79, p_FWE_ = 0.004, voxels = 47, x/y/z: 12/24/-6) and examining the overlap between the activity maps indicated that the VS and the mOFC were specifically recruited when emotional context was accounted for (**Figure 3C**). Post-hoc examination of the extracted parameter estimates revealed that activation in the posterior IFG/AI increased less during inhibition in the positive context compared to both, neutral and negative contexts (negative, left IFG/AI: p < 0.001; right IFG/AI: p = 0.033; neutral, left IFG/AI: t(226) = 5.89, p < 0.001; right IFG/AI: t(226) = 8.46, p < 0.001), whereas activation in the bilateral VS and mOFC was significantly suppressed in the neutral context as compared to both positive and negative context (positive, left VS: p < 0.001; right VS: p < 0.001; left mOFC: p < 0.001; right mOFC: p < 0.001; negative, left VS: p < 0.001; right VS: p < 0.001; left mOFC: p < 0.001; right mOFC: p < 0.001, **Figure 3D**).

**Table 1.**
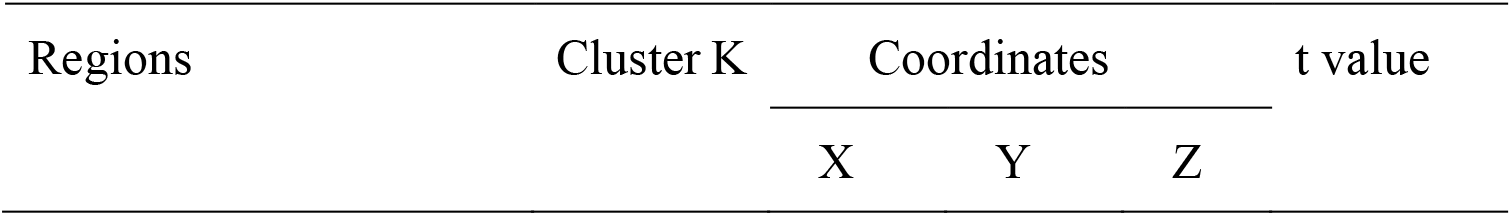
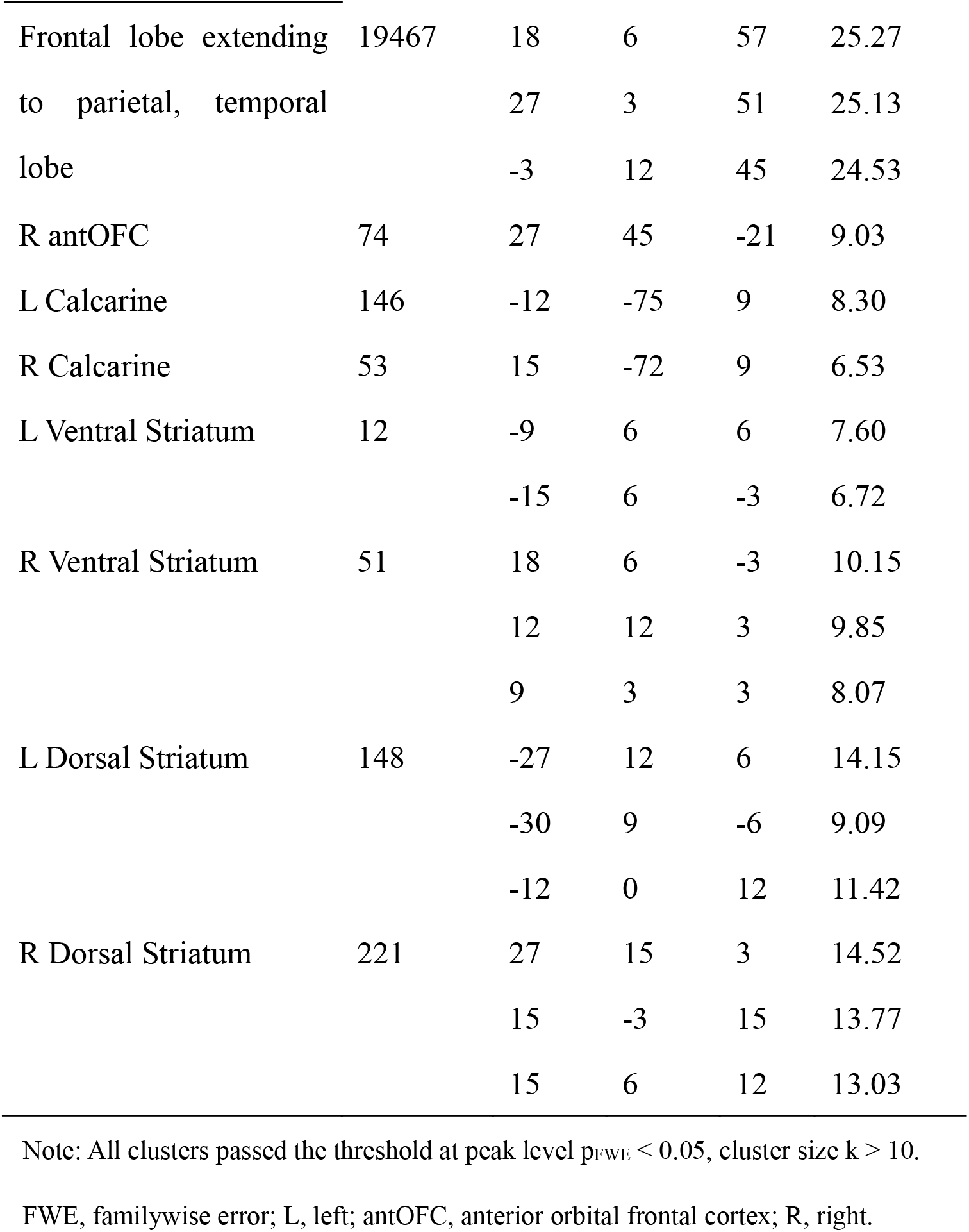
Brain regions showing general Go/No Go network on whole brain level.

**Figure 3.**
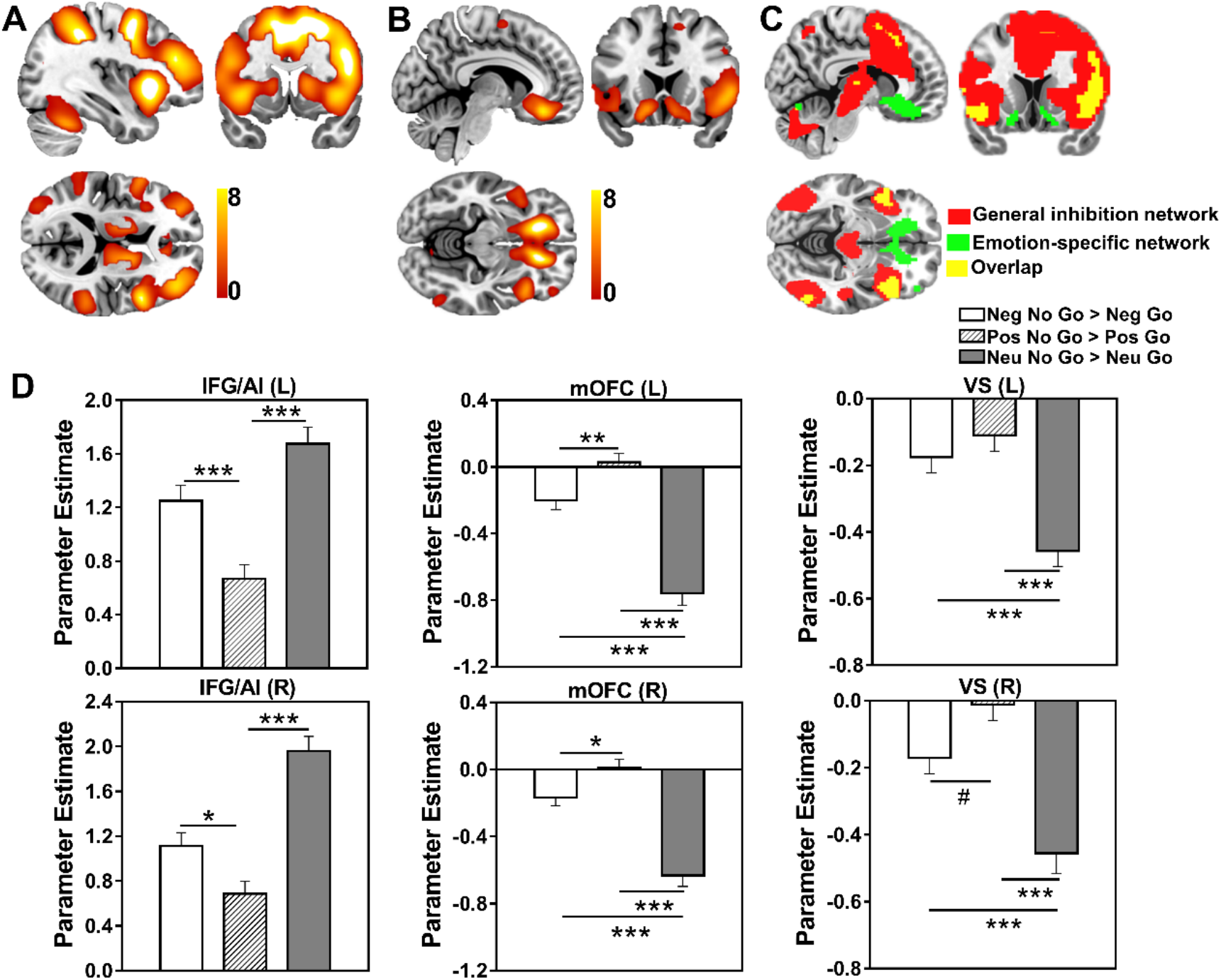
Brain activation patterns during general inhibition and interactions with emotional context. (A) General Go/No Go inhibitory network on whole brain level. (B) Activations from the interaction effect of inhibition and emotional context (contrasts: neu No Go > neu Go, neg No Go > neg Go, pos No Go > pos Go) on whole brain level. (C) Distribution in general Go/No Go inhibitory network and emotion-inhibition interaction network and (D) corresponding bar graphs based on extracted parameter estimates for the activated regions of interest (left IFG/AI and right IFG/AI, left mOFC and right mOFC, left VS and right VS). Bar graphs show mean ± SEM. #, * and *** denote significant post-hoc difference at p < 0.05, p_Bonferroni_ < 0.05 and p_Bonferroni_ < 0.001 respectively.

#### 3.2.2 Follow-up functional connectivity analysis

To further determine networks that couple in a general versus emotion-specific fashion with the identified VS regions a follow-up seed-to-voxel connectivity analysis with the left and right VS as determined by the BOLD level analysis was conducted. The left VS exhibited general negative association with fronto-parietal, temporal and limbic regions during inhibition (contrasts: No Go > Go, **Figure 4A** and **Table 2**), while the examination of the interaction between emotion and inhibition (contrasts: neu No Go > neu Go, neg No Go > neg Go, pos No Go > pos Go) revealed interactive effects on left VS coupling with the right IFG/insula (right IFG/insula, Z = 4.49, p_FWE_ < 0.001, x/y/z: 30/27/3, **Figure 4B**). Post-hoc analyses on the extracted parameter estimates revealed that left VS connectivity with right IFG/insula was significantly decreased in the positive compared to the negative context (right IFG/insula: p = 0.016) and further decreased in the neutral context (right IFG/insula: p = 0.005, **Figure 4B**).

**Table 2.** Brain regions showing significant activations in No Go vs Go conditions on connectivity level.

| Regions | Cluster K | Coordinates |  |  | t value |
| --- | --- | --- | --- | --- | --- |
|  |  | X | Y | Z |  |
| Frontal lobe extending | 18746 | -21 | 39 | 24 | 8.05 |
| to parietal lobe |  | -6 | 18 | 36 | 7.00 |
|  |  | -39 | 9 | 21 | 6.79 |

Note: All clusters passed the threshold at cluster level $p_{FWE} < 0.05$ . FWE, familywise
error.

**Figure 4.**
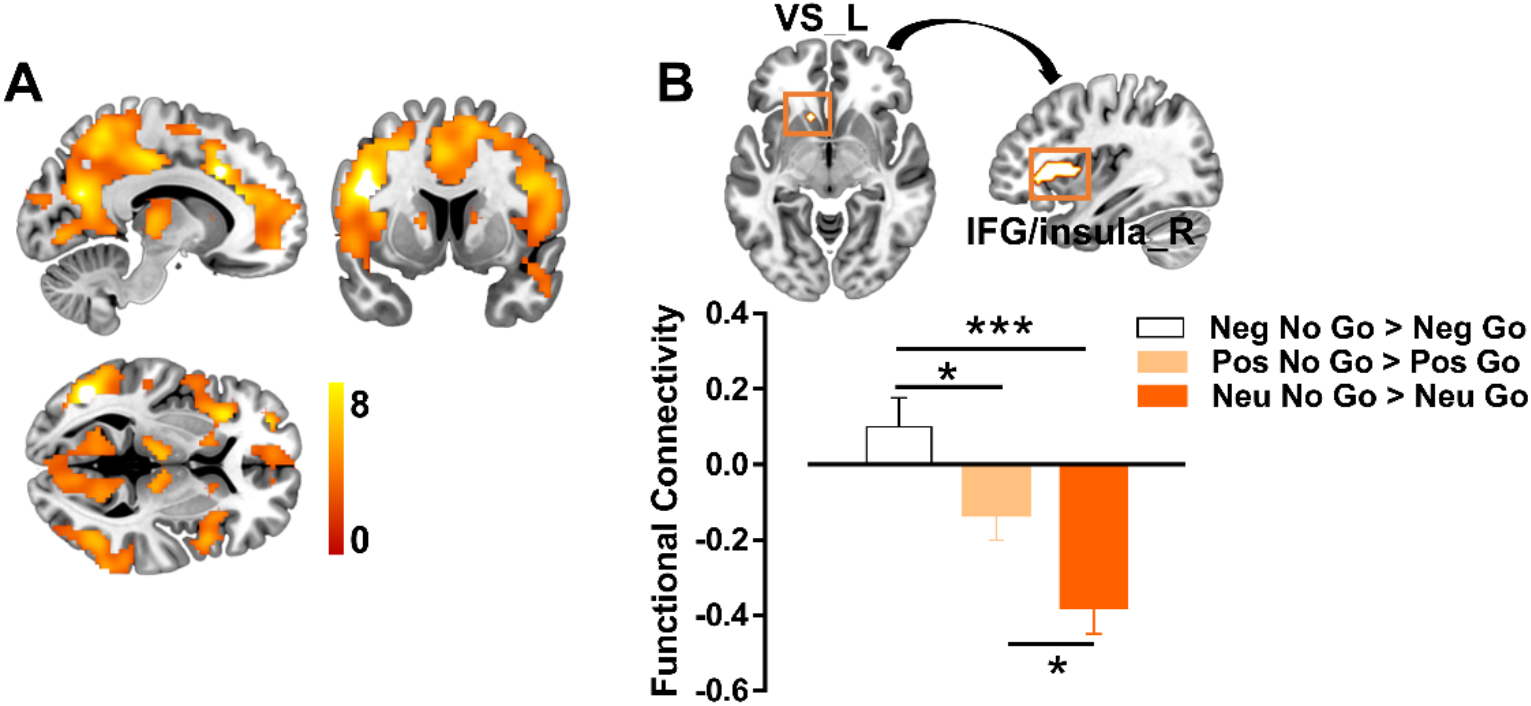
Domain-general and emotion specific functional connectivity with ventral striatum as seed. (A) Brain regions showing significant activations in No Go vs Go conditions on connectivity level. (B) Activations from the interaction effect of inhibition and emotional context (contrasts: neu No Go > neu Go, neg No Go > neg Go, pos No Go > pos Go) on connectivity level and corresponding bar graphs show functional connectivity between left VS and right IFG/insula. Bar graphs show mean ± SEM. * and *** denote significant post-hoc difference at p_Bonferroni_ < 0.05 and at p_Bonferroni_ < 0.001 respectively.

### 3.2.3 ICC Analysis

Examining the general inhibitory control networks (contrasts: No Go > Go) revealed a widespread network of increased whole brain connectivity during inhibition encompassing nearly the entire dorsal and ventral striatum as well as the limbic lobe with additional clusters located in the cerebellum, middle cingulate cortex (MCC) and fronto-temporal regions (**Figure 5A** and **Table 3**). Examination of the modulatory influence of emotional context revealed significant interaction effects in the left mOFC (Z = 3.71, p_FWE_ = 0.002, x/y/z: −18/21/-15, **Figure 5B**) and additional parietal regions (**Supplementary Table S3**) on the whole brain level. Post-hoc analyses of the extracted parameter estimates revealed that connectivity of the left mOFC significantly decreased in the neutral context during inhibition compared to both negative and positive contexts (negative, left mOFC: p < 0.001; positive, left mOFC: p < 0.001, **Figure 5B**), reflecting a valence-independent emotional modulation in this region.

**Table 3.** Brain regions showing global brain connectivity on No Go vs Go conditions from ICC analysis.

| Regions | Cluster K | Coordinates |  |  | t value |
| --- | --- | --- | --- | --- | --- |
|  |  | X | Y | Z |  |
| Limbic lobe extending to cerebellum, temporal, frontal lobe | 9774 | -6 | 9 | -27 | 13.91 |
|  |  | -15 | -9 | -45 | 13.20 |
|  |  | 51 | -39 | -30 | 12.87 |
| R MCC extending to SMA | 155 | 15 | -12 | 48 | 8.10 |
|  |  | 9 | -21 | 57 | 5.08 |
|  |  | 15 | 24 | 30 | 5.47 |
| R Postcentral gyrus | 40 | 48 | -6 | 30 | 4.55 |
|  |  | 42 | -15 | 33 | 4.32 |
|  |  | 48 | -6 | 18 | 4.08 |
| L Paracentral lobule | 102 | -9 | -21 | 60 | 5.18 |
|  |  | -9 | -24 | 51 | 5.00 |
| L Dorsal Striatum | 262 | -12 | 12 | 18 | 7.52 |
|  |  | -24 | 12 | -9 | 7.06 |
|  |  | -18 | -15 | 24 | 7.02 |
| R Dorsal Striatum | 263 | 18 | 21 | 6 | 8.94 |
|  |  | 12 | 15 | 18 | 8.56 |
|  |  | 30 | -6 | 12 | 5.42 |
| L Ventral Striatum | 202 | -15 | 0 | -9 | 8.91 |
|  |  | -9 | 6 | -15 | 7.37 |
|  |  | -21 | 9 | -9 | 6.98 |
| R Ventral Striatum | 196 | 18 | 21 | 3 | 8.12 |
|  |  | 12 | 18 | -6 | 7.10 |
|  |  | 6 | 9 | 0 | 6.95 |

Note: All clusters passed the threshold at cluster level $p_{FWE} < 0.05$ . FWE, familywise error; L, left; MCC, middle cingulum cortex; MTG, middle temporal gyrus; R, right; SFG, superior frontal gyrus; SMA, supplementary motor area.

**Figure 5.**
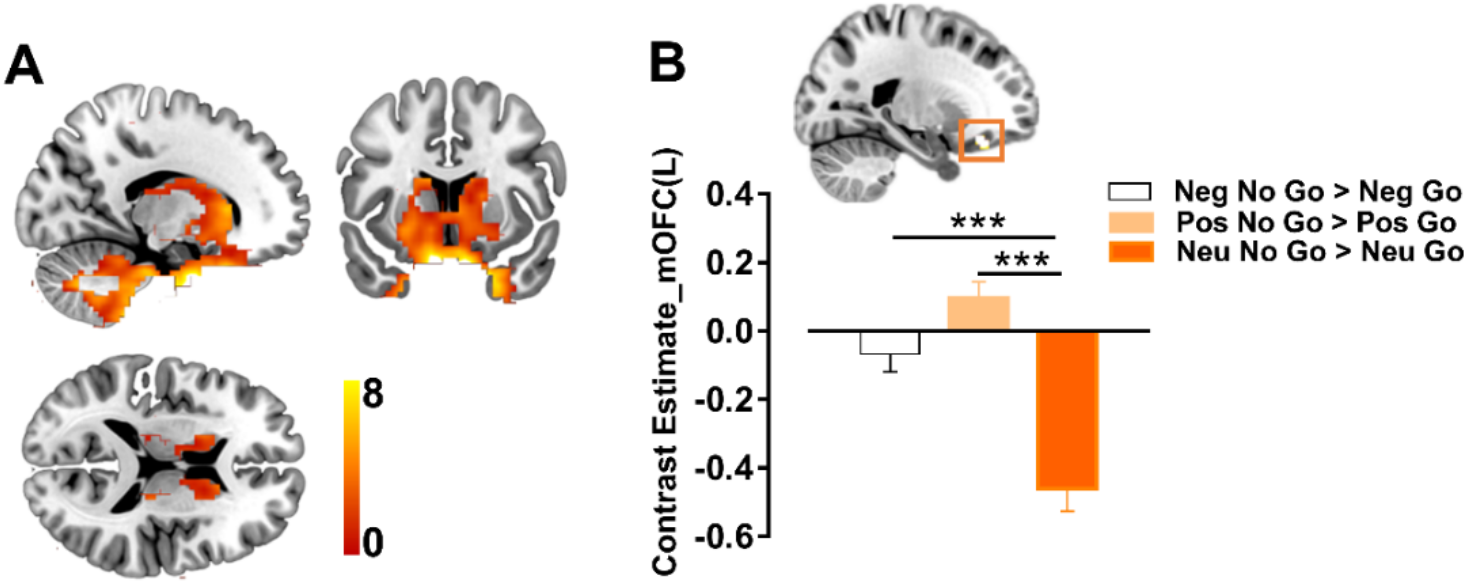
Brain regions showed global brain connectivity in general inhibition and emotion specific inhibition. (A) Brain regions showing global brain connectivity on No Go vs Go conditions from ICC analysis. (B) Emotion’s modulatory effects (contrasts: neu No Go > neu Go, neg No Go > neg Go, pos No Go > pos Go) showed higher changes of left mOFC in global brain connectivity and corresponding bar graphs based on extracted parameter estimates. Bar graphs show mean ± SEM. *** denote significant post-hoc difference at p_Bonferroni_ < 0.001.

#### 3.2.4 Follow-up Functional Connectivity Analysis

To further determine the specific fronto-striatal circuits that exhibit an emotional modulation during inhibitory control a follow-up seed-to-voxel functional connectivity analysis of the identified left mOFC region was conducted. The analysis revealed that this region exhibited a generally (emotion-unspecific, contrasts: No Go > Go) negative correlation with a widespread fronto-parietal, occipital and temporal regions during inhibition (**Figure 6A** and **Table 4**). Interaction effects of emotion on inhibition-related functional connectivity of the left mOFC were observed within the striatum (SVC corrected for the entire striatum), primarily in the left ventral part (Z = 3.81, p_FWE_ = 0.006 SVC for the entire striatum, voxels = 22, x/y/z: − 15/6/-12, **Figure 6B and Table S4**). Post-hoc analysis on extracted parameter estimates showed that mOFC connectivity with the left VS was decreased for the neutral as compared to both positive and negative contexts (positive: p = 0.004; negative: p < 0.001, **Figure 6B**).

**Table 4.**
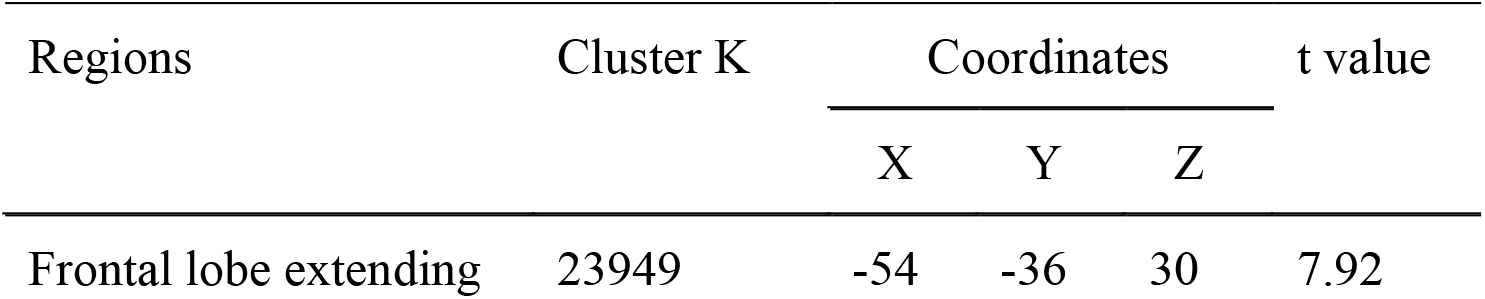
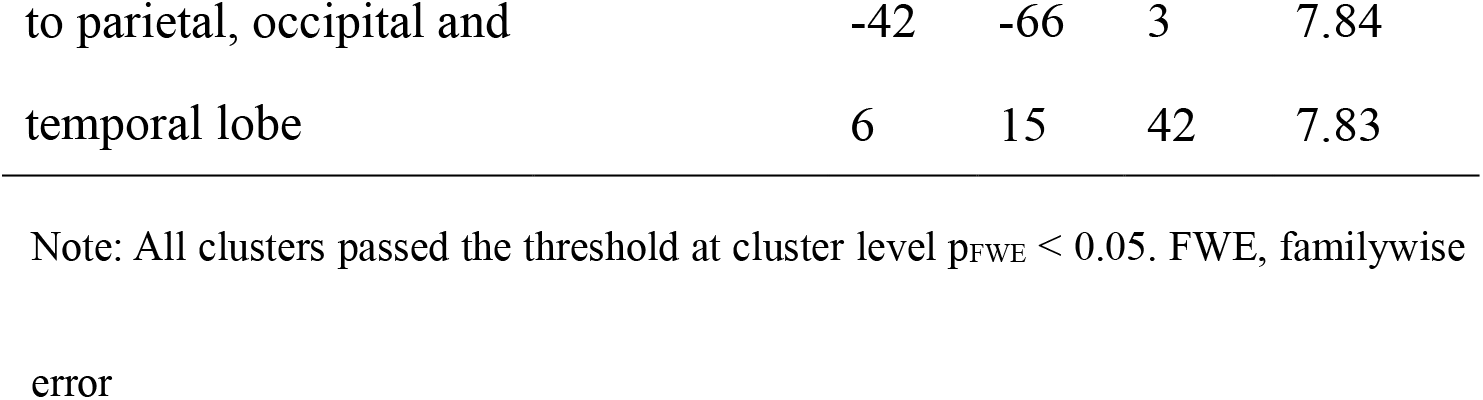
Brain regions showing significant activations in No Go vs Go condition on connectivity level.

**Figure 6.**
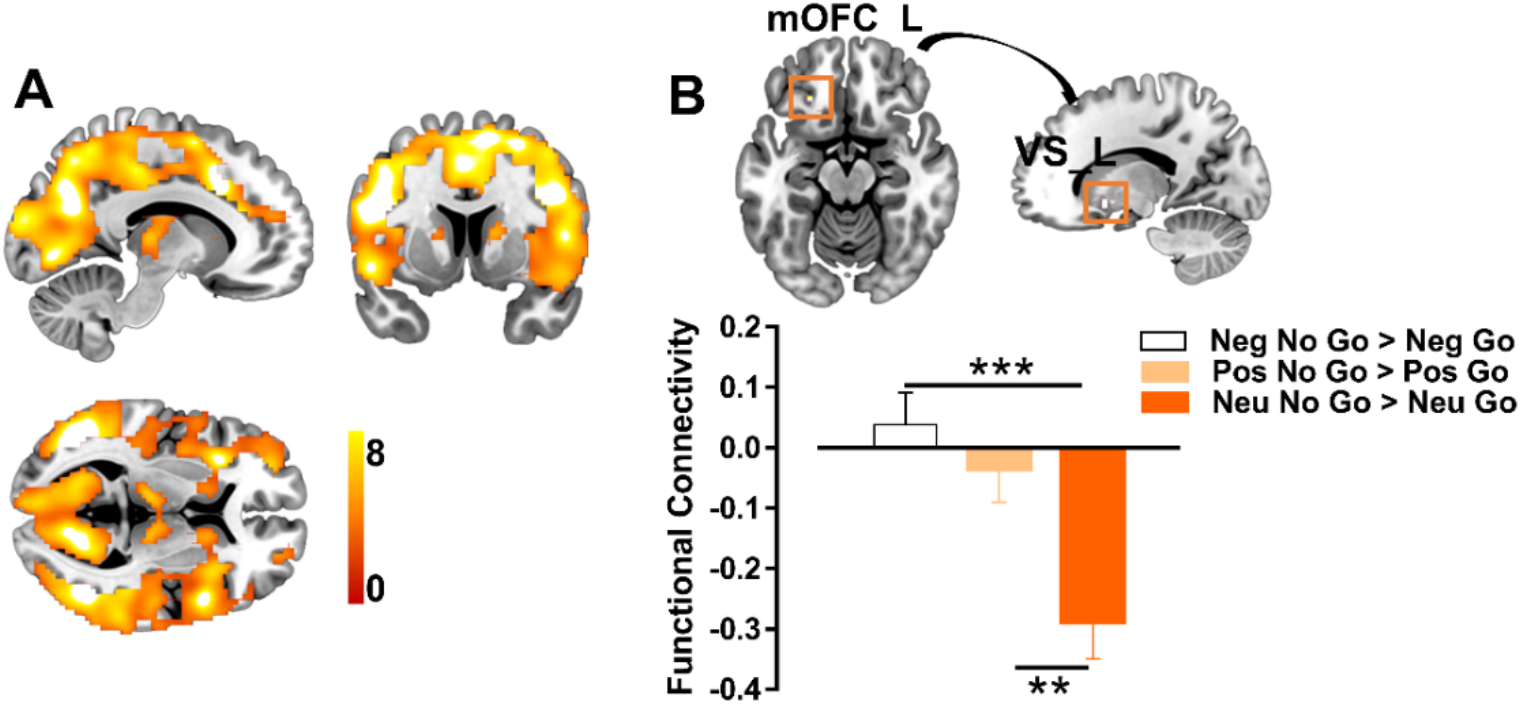
Domain-general and emotion specific functional connectivity with mOFC as seed. (A) Brain regions showing significant activations in No Go vs Go condition for connectivity results. (B) Activations from the interaction effect of inhibition and emotional context (contrasts: neu No Go > neu Go, neg No Go > neg Go, pos No Go > pos Go) on connectivity level and corresponding bar graphs show functional connectivity between left mOFC and left VS. Bar graphs show mean ± SEM, ** and *** denote significant post-hoc difference at p_Bonferroni_ < 0.01 and p_Bonferroni_ < 0.001 respectively.

## 4. Discussion

The present study aimed at segregating brain systems and circuits that mediate domain-general versus emotional context-specific inhibitory control by means of an emotional Go/No Go paradigm with concomitant fMRI acquisition in a large sample of healthy subjects. On the behavioral level more commission errors (No Go trials) and faster response times (Go trials) were observed in the positive context compared to both the negative and neutral contexts, reflecting successful emotional modulation of inhibitory control. On the neural level, regional activity in dorsal striatal, anterior insular and lateral frontal regions increased independent of emotional context while inhibition-related activity in the ventral striatum, anterior insula and medial orbitofrontal cortex varied as a function of emotional context such that the AI exhibited increased activation, while the VS and mOFC exhibited less suppression during inhibition in positive contexts. On the network level cerebellar, limbic and striatal systems generally increased voxel-wise whole brain connectivity during inhibitory control, while whole brain connectivity of the left mOFC was less suppressed during emotional contexts. Follow-up functional connectivity analyses moreover revealed that negative functional coupling between the left VS and the contralateral IFG/insula as well as the left mOFC and the ipsilateral striatum, specifically the VS, during inhibition in neutral contexts was attenuated during inhibition in both emotional contexts. Taken together, the findings indicate separable domain general as well emotional context specific inhibitory brain systems.

In line with previous studies employing emotional Go/No Go paradigms we observed higher error rates for No Go trials as compared to Go trials (Albert et al., 2010) as well as more commission errors for No Go trials in the positive context (Albert et al., 2012; Hare et al., 2005; Putman et al., 2010) as compared to both, negative and neutral contexts, reflecting successful emotional modulation of response inhibition. Together with faster response times for positive stimuli this may reflect approach tendencies for positive stimuli facilitating approach while reducing avoidance (Albert et al., 2010).

Examination of inhibition-related brain activity across emotional contexts revealed increased activity in a broad bilateral fronto-parietal cognitive control network, encompassing the lateral frontal, anterior insular, supplementary motor and superior parietal cortices, as well as striatal, particularly dorsal striatal regions. The involvement of the fronto-parietal cognitive control network is in line with previous neuroimaging studies in humans suggesting a general involvement of the fronto-parietal cognitive control network across executive domains (Chambers et al., 2009; Hung et al., 2018; Langner et al., 2018), while the engagement of primary dorsal striatal regions is in line with animal models suggesting a critical role of these regions in cognitive and motor control (Eagle et al., 2008, 2011; Eagle and Baunez, 2010; Haber, 2016).

Notably, interactions between emotional context and inhibitory control specifically engaged more medial prefrontal, particularly medial orbitofrontal, and ventral regions of the striatum. While animal and human studies demonstrated a critical role of the DS in cognitive and motor inhibitory control (Eagle et al., 2010, 2011; Ghahremani et al., 2012; Robertson et al., 2015) the VS and mOFC appear not critical for response inhibition per se (Robertson et al., 2015; Stalnaker et al., 2015) but orchestrate several emotional and reward-related processes, such that previous meta-analyses of human neuroimaging studies revealed convergent activation during positive and negative valence processing, as well as reward and decision making (De La Vega et al., 2016; Diekhof and Gruber, 2010; Hiser and Koenigs, 2018; Yang et al., 2020). Examination of extracted parameter estimates revealed that both regions were suppressed during inhibitory control in neutral contexts, while suppression was attenuated or abolished in emotional contexts, suggesting that the task-irrelevant emotional information interfered with this mechanism and may reflect the involvement of the VS-mOFC system in detecting biologically significant stimuli (Cooch et al., 2015). In contrast to our hypotheses no emotional context-sensitive interactions with inhibitory control were observed in the amygdala. This is in line with previous inhibitory control studies employing emotional linguistic stimuli (Goldstein et al., 2007), while other studies employing human face stimuli reported that amygdala activity varied as a function of emotional context (Hare et al., 2005). Given the pivotal role of the amygdala in social processes, including facial emotion recognition (Becker et al., 2012), the divergence may point to a specific role of the amygdala in inhibitory control in social contexts.

The second main aim of the study was to determine general and emotional-context specific changes on the network level during inhibitory control. Examination of voxel-level whole brain connectivity profiles revealed generally increased global connectivity of subcortical systems, including cerebellar, limbic and striatal, regions during inhibitory control while interactions between inhibitory control and emotional context specifically modulated connectivity profiles in the left precuneus and left mOFC, such that network level communication of these regions was suppressed during inhibitory control in neutral contexts yet less suppressed during negative or enhanced during positive contexts. While conceptual network-level approaches to inhibitory control emphasize the role of fronto-striatal cicruits in general inhibitory motor control (Alexander et al., 1986; Jahanshahi et al., 2015), more recent animal and human studies indicate that a neural network comprising mOFC and ventral striatal and limbic regions may mediate emotion inhibition interactions and reward related impulsive control (Goldstein et al., 2007; Hampton et al., 2017; Schulz et al., 2014; Wang et al., 2019).

Specifically, animal and human lesion studies have demonstrated the critical role of the mOFC in emotion processing as well as goal-directed response execution (Gourley et al., 2010; Mar et al., 2011; Noonan et al., 2010; Pujara et al., 2016; Shamay-Tsoory et al., 2003). In line with these findings, the mOFC changes during inhibitory control in emotional contexts emphasize the role of this region as an integrative hub at the intersection of emotional processes and inhibitory control.

Subsequent functional connectivity analyses of both, the VS and the mOFC region exhibiting emotional interactions in the BOLD activation or ICC analyses, respectively, suggest an important role of the ventral striatal-prefrontal communication in mediating the effects of emotional context on inhibition. Whereas both regions exhibited a generally strong negative coupling with core regions of the fronto-parietal control network, negative coupling between the VS and IFG/insula as well as mOFC and VS in neutral contexts was attenuated in positive contexts or switched to positive coupling in negative contexts.

The domain-general connectivity pattern of the VS is consistent with previous studies reporting negative connectivity of VS with fronto-parietal regions involved in inhibitory control (Behan et al., 2015; Diekhof and Gruber, 2010; Diekhof et al., 2012). Moreover, the context-independent negative coupling between mOFC with frontal parietal regions involved in executive control, is in accordance with previous findings suggesting a competing interaction between reward processing of the mOFC regions and executive control systems, including the dorsolateral PFC as well as parietal regions with the strength of the competing interaction being associated with inhibitory control on the behavioral and trait level (McClure et al., 2004; Zhai et al., 2015).

In line with the BOLD activity findings specifically the VS exhibited emotional context dependent modulations. Previous animal models demonstrated that - in line with focal VS lesions – VS-prefrontal disconnections produce marked impairments in the influence of emotional context on inhibitory control (Christakou et al., 2004; Feja and Koch, 2015). Given that previous animal and human studies examining structural connection between prefrontal/insular and ventral striatal regions suggest an important role in social decision making (Chikama et al., 1997; Leong et al., 2016; Rogers-Carter et al., 2019), as well as interoceptive sensitivity and behavioral control (Jaramillo et al., 2018), enhanced PFC-VS in negative relative to positive contexts may reflect a higher salience of negative stimuli during inhibitory control.

There are several limitations in the current study. First, while only fear, happy and neutral word stimuli were used in the present study, sad or angry word stimuli were not included and whether they may have differential influence on inhibition control should be investiagated in future studies. Second, only young adults were enrolled in the present study. Given that previous studies have revelaed an age-dependent effects on inhibition control (Somerville et al., 2011), it is possible that our current findings could not extend to individuals in other age groups.

In conclusion, using a validated emotional Go/No Go task in a large cohort of subjects (N = 250) allowed us to separate brain systems and specific circuits that mediated domain-general and emotional context-specific inhibitory control. The findings suggest a specific role of the VS and its connections with mOFC and insular regions in mediating the impact of emotional context on inhibitory control. The findings moreover suggest that emotional contexts - especially positive contexts – may interfere with inhibitory control system. The findings may have moreover implications for substance addiction as well as neuropsychiatric disorders characterized by deficient inhibitory control and enhanced reward sensitivity.

## Acknowledgements

This work was supported by the National Key Research and Development Program of China (Grant No. 2018YFA0701400), National Natural Science Foundation of China (NSFC No. 31700998, 31530032).

## References

Albert, J., López-Martín, S., Carretié, L., 2010. Emotional context modulates response inhibition: neural and behavioral data. Neuroimage 49, 914–921.

Albert, J., López-Martín, S., Tapia, M., Montoya, D., Carretie, L., 2012. The role of the anterior cingulate cortex in emotional response inhibition. Human brain mapping 33, 2147–2160.

Alexander, G.E., DeLong, M.R., Strick, P.L., 1986. Parallel organization of functionally segregated circuits linking basal ganglia and cortex. Annual review of neuroscience 9, 357–381.

Aron, A.R., Monsell, S., Sahakian, B.J., Robbins, T.W., 2004. A componential analysis of task-switching deficits associated with lesions of left and right frontal cortex. Brain 127, 1561–1573.

Becker, B., Mihov, Y., Scheele, D., Kendrick, K.M., Feinstein, J.S., Matusch, A., Aydin, M., Reich, H., Urbach, H., Oros-Peusquens, A.-M., 2012. Fear processing and social networking in the absence of a functional amygdala. Biological psychiatry 72, 70–77.

Behan, B., Stone, A., Garavan, H., 2015. Right prefrontal and ventral striatum interactions underlying impulsive choice and impulsive responding. Human brain mapping 36, 187–198.

Berkman, E.T., Burklund, L., Lieberman, M.D., 2009. Inhibitory spillover: Intentional motor inhibition produces incidental limbic inhibition via right inferior frontal cortex. Neuroimage 47, 705–712.

Brown, M.R., Lebel, R.M., Dolcos, F., Wilman, A.H., Silverstone, P.H., Pazderka, H., Fujiwara, E., Wild, T.C., Carroll, A.M., Hodlevskyy, O., 2012. Effects of emotional context on impulse control. Neuroimage 63, 434–446.

Chambers, C.D., Garavan, H., Bellgrove, M.A., 2009. Insights into the neural basis of response inhibition from cognitive and clinical neuroscience. Neuroscience & Biobehavioral Reviews 33, 631–646.

Chang, J., Hu, J., Li, C.-S.R., Yu, R., 2020. Neural correlates of enhanced response inhibition in the aftermath of stress. Neuroimage 204, 116212.

Chen, T., Becker, B., Camilleri, J., Wang, L., Yu, S., Eickhoff, S.B., Feng, C., 2018. A domain-general brain network underlying emotional and cognitive interference processing: evidence from coordinate-based and functional connectivity meta-analyses. Brain Structure and Function 223, 3813–3840.

Chikama, M., McFarland, N.R., Amaral, D.G., Haber, S.N., 1997. Insular cortical projections to functional regions of the striatum correlate with cortical cytoarchitectonic organization in the primate. Journal of Neuroscience 17, 9686–9705.

Christakou, A., Robbins, T.W., Everitt, B.J., 2004. Prefrontal cortical–ventral striatal interactions involved in affective modulation of attentional performance: implications for corticostriatal circuit function. Journal of Neuroscience 24, 773–780.

Cooch, N.K., Stalnaker, T.A., Wied, H.M., Bali-Chaudhary, S., McDannald, M.A., Liu, T.-L., Schoenbaum, G., 2015. Orbitofrontal lesions eliminate signalling of biological significance in cue-responsive ventral striatal neurons. Nature Communications 6, 7195.

Courtney, K.E., Ghahremani, D.G., Ray, L.A., 2013. Fronto-striatal functional connectivity during response inhibition in alcohol dependence. Addiction biology 18, 593–604.

Cromheeke, S., Mueller, S.C., 2014. Probing emotional influences on cognitive control: an ALE meta-analysis of cognition emotion interactions. Brain Structure and Function 219, 995–1008.

Dalley, J.W., Robbins, T.W., 2017. Fractionating impulsivity: neuropsychiatric implications. Nature Reviews Neuroscience 18, 158.

De La Vega, A., Chang, L.J., Banich, M.T., Wager, T.D., Yarkoni, T., 2016. Large-scale meta-analysis of human medial frontal cortex reveals tripartite functional organization. Journal of Neuroscience 36, 6553–6562.

Diekhof, E.K., Gruber, O., 2010. When desire collides with reason: functional interactions between anteroventral prefrontal cortex and nucleus accumbens underlie the human ability to resist impulsive desires. Journal of Neuroscience 30, 1488–1493.

Diekhof, E.K., Nerenberg, L., Falkai, P., Dechent, P., Baudewig, J., Gruber, O., 2012. Impulsive personality and the ability to resist immediate reward: An fMRI study examining interindividual differences in the neural mechanisms underlying self-control. Human brain mapping 33, 2768–2784.

Diener, C., Kuehner, C., Brusniak, W., Ubl, B., Wessa, M., Flor, H., 2012. A meta-analysis of neurofunctional imaging studies of emotion and cognition in major depression. Neuroimage 61, 677–685.

Drevets, W.C., Raichle, M.E., 1998. Reciprocal suppression of regional cerebral blood flow during emotional versus higher cognitive processes: Implications for interactions between emotion and cognition. Cognition and emotion 12, 353–385.

Eagle, D.M., Bari, A., Robbins, T.W., 2008. The neuropsychopharmacology of action inhibition: cross-species translation of the stop-signal and go/no-go tasks. Psychopharmacology 199, 439–456.

Eagle, D.M., Baunez, C., 2010. Is there an inhibitory-response-control system in the rat? Evidence from anatomical and pharmacological studies of behavioral inhibition. Neuroscience & Biobehavioral Reviews 34, 50–72.

Eagle, D.M., Wong, J.C., Allan, M.E., Mar, A.C., Theobald, D.E., Robbins, T.W., 2011. Contrasting roles for dopamine D1 and D2 receptor subtypes in the dorsomedial striatum but not the nucleus accumbens core during behavioral inhibition in the stop-signal task in rats. Journal of Neuroscience 31, 7349–7356.

Fan, L., Li, H., Zhuo, J., Zhang, Y., Wang, J., Chen, L., Yang, Z., Chu, C., Xie, S., Laird, A.R., 2016. The human brainnetome atlas: a new brain atlas based on connectional architecture. Cerebral cortex 26, 3508–3526.

Feja, M., Koch, M., 2015. Frontostriatal systems comprising connections between ventral medial prefrontal cortex and nucleus accumbens subregions differentially regulate motor impulse control in rats. Psychopharmacology 232, 1291–1302.

Ghahremani, D.G., Lee, B., Robertson, C.L., Tabibnia, G., Morgan, A.T., De Shetler, N., Brown, A.K., Monterosso, J.R., Aron, A.R., Mandelkern, M.A., 2012. Striatal dopamine D2/D3 receptors mediate response inhibition and related activity in frontostriatal neural circuitry in humans. Journal of Neuroscience 32, 7316–7324.

Goldstein, M., Brendel, G., Tuescher, O., Pan, H., Epstein, J., Beutel, M., Yang, Y., Thomas, K., Levy, K., Silverman, M., 2007. Neural substrates of the interaction of emotional stimulus processing and motor inhibitory control: an emotional linguistic go/no-go fMRI study. Neuroimage 36, 1026–1040.

Gourley, S.L., Lee, A.S., Howell, J.L., Pittenger, C., Taylor, J.R., 2010. Dissociable regulation of instrumental action within mouse prefrontal cortex. European Journal of Neuroscience 32, 1726–1734.

Haber, S.N., 2016. Corticostriatal circuitry. Dialogues in clinical neuroscience 18, 7.

Hampton, W.H., Alm, K.H., Venkatraman, V., Nugiel, T., Olson, I.R., 2017. Dissociable frontostriatal white matter connectivity underlies reward and motor impulsivity. Neuroimage 150, 336–343.

Hare, T.A., Tottenham, N., Davidson, M.C., Glover, G.H., Casey, B., 2005. Contributions of amygdala and striatal activity in emotion regulation. Biological psychiatry 57, 624–632.

Hiser, J., Koenigs, M., 2018. The multifaceted role of the ventromedial prefrontal cortex in emotion, decision making, social cognition, and psychopathology. Biological psychiatry 83, 638–647.

Hu, Y., Salmeron, B.J., Krasnova, I.N., Gu, H., Lu, H., Bonci, A., Cadet, J.L., Stein, E.A., Yang, Y., 2019. Compulsive drug use is associated with imbalance of orbitofrontal-and prelimbic-striatal circuits in punishment-resistant individuals. Proceedings of the National Academy of Sciences 116, 9066–9071.

Hung, Y., Gaillard, S.L., Yarmak, P., Arsalidou, M., 2018. Dissociations of cognitive inhibition, response inhibition, and emotional interference: Voxelwise ALE meta-analyses of fMRI studies. Human brain mapping 39, 4065–4082.

Jahanshahi, M., Obeso, I., Rothwell, J.C., Obeso, J.A., 2015. A fronto–striato–subthalamic– pallidal network for goal-directed and habitual inhibition. Nature Reviews Neuroscience 16, 719–732.

Jaramillo, A.A., Van Voorhies, K., Randall, P.A., Besheer, J., 2018. Silencing the insular-striatal circuit decreases alcohol self-administration and increases sensitivity to alcohol. Behavioural brain research 348, 74–81.

Karalunas, S.L., Weigard, A., Alperin, B., 2020. Emotion-Cognition Interactions in Attention-Deficit/Hyperactivity Disorder: Increased Early Attention Capture and Weakened Attentional Control in Emotional Contexts. Biological psychiatry. Cognitive Neuroscience and Neuroimaging.

Kelly, A.C., Hester, R., Murphy, K., Javitt, D.C., Foxe, J.J., Garavan, H., 2004. Prefrontal-subcortical dissociations underlying inhibitory control revealed by event-related fMRI. European Journal of Neuroscience 19, 3105–3112.

Lüscher, C., Robbins, T.W., Everitt, B.J., 2020. The transition to compulsion in addiction. Nature Reviews Neuroscience, 1–17.

Langner, R., Leiberg, S., Hoffstaedter, F., Eickhoff, S.B., 2018. Towards a human self-regulation system: common and distinct neural signatures of emotional and behavioural control. Neuroscience & Biobehavioral Reviews 90, 400–410.

Leong, J.K., Pestilli, F., Wu, C.C., Samanez-Larkin, G.R., Knutson, B., 2016. White-matter tract connecting anterior insula to nucleus accumbens correlates with reduced preference for positively skewed gambles. Neuron 89, 63–69.

Li, C.-s.R., Sinha, R., 2008. Inhibitory control and emotional stress regulation: Neuroimaging evidence for frontal–limbic dysfunction in psycho-stimulant addiction. Neuroscience & Biobehavioral Reviews 32, 581–597.

Liu, C., Dai, J., Chen, Y., Qi, Z., Xin, F., Zhuang, Q., Zhou, X., Zhou, F., Luo, L., Huang, Y., 2020. Disorder-and emotional context-specific neurofunctional alterations during inhibitory control in generalized anxiety disorder and major depressive disorder.

Mar, A.C., Walker, A.L., Theobald, D.E., Eagle, D.M., Robbins, T.W., 2011. Dissociable effects of lesions to orbitofrontal cortex subregions on impulsive choice in the rat. Journal of Neuroscience 31, 6398–6404.

Martuzzi, R., Ramani, R., Qiu, M., Shen, X., Papademetris, X., Constable, R.T., 2011. A whole-brain voxel based measure of intrinsic connectivity contrast reveals local changes in tissue connectivity with anesthetic without a priori assumptions on thresholds or regions of interest. Neuroimage 58, 1044–1050.

McClure, S.M., Laibson, D.I., Loewenstein, G., Cohen, J.D., 2004. Separate neural systems value immediate and delayed monetary rewards. Science 306, 503–507.

McLaren, D.G., Ries, M.L., Xu, G., Johnson, S.C., 2012. A generalized form of context-dependent psychophysiological interactions (gPPI): a comparison to standard approaches. Neuroimage 61, 1277–1286.

Meffert, H., Hwang, S., Nolan, Z.T., Chen, G., Blair, J.R., 2016. Segregating attention from response control when performing a motor inhibition task: segregating attention from response control. Neuroimage 126, 27–38.

Morein-Zamir, S., Robbins, T.W., 2015. Fronto-striatal circuits in response-inhibition: Relevance to addiction. Brain research 1628, 117–129.

Noonan, M., Walton, M., Behrens, T., Sallet, J., Buckley, M., Rushworth, M., 2010. Separate value comparison and learning mechanisms in macaque medial and lateral orbitofrontal cortex. Proceedings of the National Academy of Sciences 107, 20547–20552.

Protopopescu, X., Pan, H., Altemus, M., Tuescher, O., Polanecsky, M., McEwen, B., Silbersweig, D., Stern, E., 2005. Orbitofrontal cortex activity related to emotional processing changes across the menstrual cycle. Proceedings of the National Academy of Sciences 102, 16060–16065.

Pujara, M.S., Philippi, C.L., Motzkin, J.C., Baskaya, M.K., Koenigs, M., 2016. Ventromedial prefrontal cortex damage is associated with decreased ventral striatum volume and response to reward. Journal of Neuroscience 36, 5047–5054.

Putman, P., van Peer, J., Maimari, I., van der Werff, S., 2010. EEG theta/beta ratio in relation to fear-modulated response-inhibition, attentional control, and affective traits. Biological psychology 83, 73–78.

Roberts, G., Green, M.J., Breakspear, M., McCormack, C., Frankland, A., Wright, A., Levy, F., Lenroot, R., Chan, H.N., Mitchell, P.B., 2013. Reduced inferior frontal gyrus activation during response inhibition to emotional stimuli in youth at high risk of bipolar disorder. Biological psychiatry 74, 55–61.

Robertson, C.L., Ishibashi, K., Mandelkern, M.A., Brown, A.K., Ghahremani, D.G., Sabb, F., Bilder, R., Cannon, T., Borg, J., London, E.D., 2015. Striatal D1-and D2-type dopamine receptors are linked to motor response inhibition in human subjects. Journal of Neuroscience 35, 5990–5997.

Rogers-Carter, M.M., Djerdjaj, A., Gribbons, K.B., Varela, J.A., Christianson, J.P., 2019. Insular cortex projections to nucleus accumbens core mediate social approach to stressed juvenile rats. Journal of Neuroscience 39, 8717–8729.

Schulz, K.P., Bédard, A.-C.V., Fan, J., Clerkin, S.M., Dima, D., Newcorn, J.H., Halperin, J.M., 2014. Emotional bias of cognitive control in adults with childhood attention-deficit/hyperactivity disorder. NeuroImage: Clinical 5, 1–9.

Schulz, K.P., Fan, J., Magidina, O., Marks, D.J., Hahn, B., Halperin, J.M., 2007. Does the emotional go/no-go task really measure behavioral inhibition? Convergence with measures on a non-emotional analog. Archives of Clinical Neuropsychology 22, 151–160.

Shafritz, K.M., Collins, S.H., Blumberg, H.P., 2006. The interaction of emotional and cognitive neural systems in emotionally guided response inhibition. Neuroimage 31, 468–475.

Shamay-Tsoory, S.G., Tomer, R., Berger, B.D., Aharon-Peretz, J., 2003. Characterization of empathy deficits following prefrontal brain damage: the role of the right ventromedial prefrontal cortex. Journal of cognitive neuroscience 15, 324–337.

Somerville, L.H., Hare, T., Casey, B., 2011. Frontostriatal maturation predicts cognitive control failure to appetitive cues in adolescents. Journal of cognitive neuroscience 23, 2123–2134.

Stalnaker, T.A., Cooch, N.K., Schoenbaum, G., 2015. What the orbitofrontal cortex does not do. Nature neuroscience 18, 620.

Swick, D., Ashley, V., Turken, U., 2008. Left inferior frontal gyrus is critical for response inhibition. BMC neuroscience 9, 1–11.

Taylor, M.J., Robertson, A., Keller, A.E., Sato, J., Urbain, C., Pang, E.W., 2018. Inhibition in the face of emotion: Characterization of the spatial-temporal dynamics that facilitate automatic emotion regulation. Human brain mapping 39, 2907–2916.

Todd, R.M., Lee, W., Evans, J.W., Lewis, M.D., Taylor, M.J., 2012. Withholding response in the face of a smile: age-related differences in prefrontal sensitivity to Nogo cues following happy and angry faces. Developmental cognitive neuroscience 2, 340–350.

Torrisi, S., Robinson, O., O’Connell, K., Davis, A., Balderston, N., Ernst, M., Grillon, C., 2016. The neural basis of improved cognitive performance by threat of shock. Social Cognitive and Affective Neuroscience 11, 1677–1686.

Tottenham, N., Hare, T. A., Millner, A., Gilhooly, T., Zevin, J. D., Casey, B. J. 2011. Elevated amygdala response to faces following early deprivation. Developmental science 14, 190–204.

Tsvetanov, K.A., Ye, Z., Hughes, L., Samu, D., Treder, M.S., Wolpe, N., Tyler, L.K., Rowe, J.B., 2018. Activity and connectivity differences underlying inhibitory control across the adult life span. Journal of Neuroscience 38, 7887–7900.

Ursu, S., Kring, A.M., Gard, M.G., Minzenberg, M.J., Yoon, J.H., Ragland, J.D., Solomon, M., Carter, C.S., 2011. Prefrontal cortical deficits and impaired cognition-emotion interactions in schizophrenia. American Journal of Psychiatry 168, 276–285.

Vatansever, D., Manktelow, A., Sahakian, B., Menon, D., Stamatakis, E.A., 2017. Angular default mode network connectivity across working memory load. Human brain mapping 38, 41–52.

Vatansever, D., Menon, D.K., Manktelow, A.E., Sahakian, B.J., Stamatakis, E.A., 2015. Default mode dynamics for global functional integration. Journal of Neuroscience 35, 15254–15262.

Vercammen, A., Morris, R., Green, M.J., Lenroot, R., Kulkarni, J., Carr, V.J., Weickert, C.S., Weickert, T.W., 2012. Reduced neural activity of the prefrontal cognitive control circuitry during response inhibition to negative words in people with schizophrenia. Journal of psychiatry & neuroscience: JPN 37, 379.

Volkow, N.D., Wang, G.-J., Fowler, J.S., Tomasi, D., Telang, F., 2011. Addiction: beyond dopamine reward circuitry. Proceedings of the National Academy of Sciences 108, 15037–15042.

Wang, Z., Yue, L., Cui, C., Liu, S., Wang, X., Li, Y., Ma, L., 2019. Top-down control of the medial orbitofrontal cortex to nucleus accumbens core pathway in decisional impulsivity. Brain Structure and Function 224, 2437–2452.

Wessa, M., Houenou, J., Paillère-Martinot, M.-L., Berthoz, S., Artiges, E., Leboyer, M., Martinot, J.-L., 2007. Fronto-striatal overactivation in euthymic bipolar patients during an emotional go/nogo task. American Journal of Psychiatry 164, 638–646.

Whitfield-Gabrieli, S., Nieto-Castanon, A., 2012. Conn: a functional connectivity toolbox for correlated and anticorrelated brain networks. Brain connectivity 2, 125–141.

Xu, M., Xu, G., Yang, Y., 2016. Neural systems underlying emotional and non-emotional interference processing: An ALE meta-analysis of functional neuroimaging studies. Frontiers in behavioral neuroscience 10, 220.

Yang, M., Tsai, S.-J., Li, C.-S.R., 2020. Concurrent amygdalar and ventromedial prefrontal cortical responses during emotion processing: a meta-analysis of the effects of valence of emotion and passive exposure versus active regulation. Brain Structure and Function 225, 345–363.

Zhai, T., Shao, Y., Chen, G., Ye, E., Ma, L., Wang, L., Lei, Y., Chen, G., Li, W., Zou, F., 2015. Nature of functional links in valuation networks differentiates impulsive behaviors between abstinent heroin-dependent subjects and nondrug-using subjects. Neuroimage 115, 76–84.

Zhang, W., Ding, Q., Chen, N., Wei, Q., Zhao, C., Zhang, P., Li, X., Liu, Q., Li, H., 2016. The development of automatic emotion regulation in an implicit emotional Go/NoGo paradigm and the association with depressive symptoms and anhedonia during adolescence. NeuroImage: Clinical 11, 116–123.

