## Supplemental Information for "Segregating domain-general from emotional context-specific inhibitory control systems - ventral striatum and orbitofrontal cortex serve as emotion-cognition integration hubs"

### Supplementary Information

Zhuang et al.,

Contact:

**Table S1. Results from emotional valence, intensity, imagination rating as well as word frequency.**

| | Stimuli | | | | |
|---|---|---|---|---|---|
| Measurements | Positive | Negative | Neutral | F | p |
| Emotional valence rating | 94.58% ± 1.07% | 96.00% ± 1.07% | 96.21% ± 0.73% | 0.95 | 0.35 |
| Frequency | 1965.58 ± 340.04 | 1007.24 ± 446.22 | 1725.54 ± 288.05 | 1.87 | 0.16 |
| Intensity | 5.73 ± 0.17 | 6.15 ± 0.23 | | 1.47 (t) | 0.15 |
| Imagination | 5.76 ± 0.23 | 6.07 ± 0.22 | 5.70 ± 0.12 | 1.02 | 0.37 |

**Table S2. Brain regions exhibit inhibition x emotion effects on the whole brain level**

| Regions | Cluster K | Coordinates | | | Z value |
|---|---|---|---|---|---|
| | | X | Y | Z | |
| mOFC | 969 | -15 | 33 | -18 | Inf |
| | | 12 | 33 | -15 | Inf |
| | | 15 | 45 | -9 | 6.07 |
| L IFG/Insula | 235 | -39 | 15 | -12 | 6.13 |
| | | -48 | 12 | -12 | 5.95 |
| | | -60 | 12 | 6 | 5.41 |
| R IFG/Insula extending to Precentral | 979 | 51 | 18 | -3 | 7.45 |
| | | 45 | 21 | -9 | 7.34 |
| | | 45 | 6 | 42 | 6.20 |
| R MFG | 194 | 42 | 48 | 6 | 5.71 |

| | | | | | |
|---|---|---|---|---|---|
| | | 45 | 45 | -9 | 5.30 |
| | | 39 | 36 | 21 | 4.76 |
| R MTG | 125 | 54 | -66 | -3 | 6.23 |
| L STG | 143 | -54 | -36 | 21 | 6.26 |
| R STG | 63 | 60 | -33 | 18 | 5.44 |
| L Cerebellum | 196 | -30 | -63 | -27 | 6.06 |
| | | -18 | -75 | -21 | 4.87 |
| R Vermis | 68 | 6 | -69 | -15 | 5.71 |
| R IPL extending to Supra | 135 | 51 | -39 | 42 | 5.35 |
| Marginal | | 42 | -48 | 54 | 4.91 |
| L Postcentral | 11 | -63 | -18 | 21 | 4.95 |
| L MOG | 14 | -51 | -72 | 0 | 4.89 |
| L Thal | 24 | -15 | -12 | 9 | 5.23 |
| R Thal | 52 | 12 | -9 | 9 | 5.87 |
| L SMA | 36 | -9 | -3 | 63 | 5.48 |
| L Dorsal Striatum | 16 | -27 | -9 | 6 | 5.12 |
| | | -27 | -21 | 6 | 5.01 |
| L Ventral Striatum | 38 | -12 | 18 | -9 | 6.04 |
| | | -12 | 6 | -12 | 5.33 |
| | | -12 | 27 | -3 | 4.77 |
| R Ventral Striatum | 47 | 12 | 24 | -6 | 5.79 |
| | | 9 | 9 | -9 | 5.56 |

Note: All regions passed the threshold at peak level $p_{FWE} < 0.05$, cluster size $k > 10$.

FWE, familywise error; IFG, inferior frontal gyrus; IPL, inferior parietal lobule; L, left; MFG, middle frontal gyrus; mOFC, medial orbital frontal cortex; MOG, middle occipital gyrus; MTG, middle temporal gyrus; R, right; SMA, supplementary motor area; STG, superior temporal gyrus; Thal, Thalamus.

**Table S3. Inhibition x emotion effects from the ICC analysis.**

| Regions | Cluster K | Coordinates | | | Z value |
|---|---|---|---|---|---|
| | | X | Y | Z | |
| L Precuneus | 67 | -3 | -60 | 39 | 4.29 |
| | | -15 | -48 | 33 | 3.92 |
| L mOFC | 52 | -21 | 30 | -18 | 4.08 |
| | | -18 | 21 | -15 | 3.71 |

Note: All clusters passed the threshold at cluster level $p_{FWE} < 0.05$ for ICC results.

FWE, familywise error; L, left; mOFC, medial orbital frontal cortex.

**Table S4. Inhibition x emotion effects from functional connectivity following ICC analysis.**

| Regions | Cluster K | Coordinates | | | Z value |
|---|---|---|---|---|---|
| | | X | Y | Z | |
| L Ventral Striatum | 22 | -15 | 6 | -12 | 3.81 |
| R Ventral Striatum | 4 | 12 | 12 | 3 | 3.70 |

Note: All clusters passed the threshold at peak level pFWE < 0.05 small volume correction (SVC) for ICC results. FWE, familywise error; L, left; R, right.
